# Calcium (Ca2+) fluxes at Mitochondria-ER Contact Sites (MERCS) are a new target of senolysis in Therapy-Induced Senescence (TIS)

**DOI:** 10.1101/2024.03.31.587471

**Authors:** Andrea Puebla-Huerta, Hernán Huerta, Camila Quezada-Gutierez, Pablo Morgado-Cáceres, César Casanova-Canelo, Sergio Linsambarth, Osman Díaz-Rivera, José Alberto López-Domínguez, Sandra Rodríguez-López, Galdo Bustos, Eduardo Silva-Pavez, Alenka Lovy, Gabriel Quiroz, Catalina González-Seguel, Edison Salas-Huenuleo, Marcelo J. Kogan, Jordi Molgó, Armen Zakarian, José M. Villalba, Christian Gonzalez-Billault, Ulises Ahumada-Castro, J. César Cárdenas

## Abstract

- This study investigates the state of calcium (Ca^2+^) flux and Mitochondria-ER contact sites (MERCS) on Therapy-Induced Senescence (TIS).
- TIS cells-induced by Doxorubicin and Etoposide increase their MERCS contact surface but exhibit a decreased ER-mitochondria Ca^2+^ flux.
- TIS cells show decreased levels of IP3R isoforms and a decreased interaction between type 1 IP3R isoform and VDAC1.
- The ER-mitochondria Ca^2+^ flux is essential to maintain the viability of senescence cells.
- Inhibition of ER-mitochondria Ca^2+^ flux rise as a new target of senolysis *in vitro* and *in vivo*.

## MAIN BODY

Cellular senescence, in its canonical manifestation, is characterized by the cessation of the replicative capacity and the development of a distinctive altered phenotype. This phenotype encompasses modifications in the secretome, activation of tumor suppressor genes, and alterations in chromatin and genome integrity[1]. Various types of cellular senescence have been identified, including replicative senescence, oncogene-induced senescence (OIS), and therapy-induced senescence (TIS).

Across various types of cellular senescence, common features observed in senescent cells comprises the activation of tumor suppressor pathways, induction of senescence-associated β-galactosidase activity, and metabolic alterations, particularly associated to mitochondria[2]. Specifically, these alterations encompass mitochondrial network imbalance, changes in mitochondrial membrane potential and electron transport chain, and dysregulation of bioenergetic [3].

A pivotal event that dictates mitochondrial function is the influx of calcium (Ca^2+^) ions into the mitochondria. Ca^2+^ primarily permeates the mitochondria from the endoplasmic reticulum (ER) through microdomains where there exists a close proximity between mitochondria and the ER, forming quasi-synaptic junctions known as mitochondria-ER contact sites (MERCS). These MERCS typically exhibit a thickness ranging from 10 to 50 nm. Ca^2+^ release from the ER is mediated by the inositol 1,4,5-trisphosphate (InsP3) receptors (InsP3Rs) and Ryanodine Receptors (RyR), allowing Ca^2+^ to enter the mitochondria. Initially, Ca^2+^ passes through the Voltage Dependent Anion Channel (VDAC) located in the outer mitochondrial membrane before reaching the mitochondrial calcium uniporter complex (MCUC) situated in the inner mitochondrial membrane[4].

Enhanced MERCS, and increased ER-mitochondria Ca^2+^ fluxes have been documented in specific types of senescent cells[4–7]. However, the specificities regarding chemotherapy-induced senescence, herein denoted as TIS cells, remain obscure. To elucidate this, we initially established a TIS model. We used growing human diploid IMR-90 fibroblasts which were treated with two chemotherapeutic drugs, Doxorubicin [Doxo] (250 nM) and Etoposide [Eto] (40 μM) over 48 h. Ten days after both treatments the cells were harvested for experiments (**Figure 1A**). The evaluation in both group of cells with well-established senescence markers showed an arrest of cell proliferation with an increase in the expression of negative regulators of the cell cycle p16 and p21 (**Figure 1B**). Also, an increase in the SA-β-gal staining (**Figure 1C**) and an increase of several component of the SASP that included IL6, IL1α, IL1β, CXCL1 and MMP3 (**Figure 1D**) was also observed. In this firmly established TIS cell model, we investigated the proximity between ER and mitochondrial membranes using confocal microscopy as an initial approach, noting an increase in ER-mitochondria proximity in both Doxorubicin and Etoposide-induced TIS (Colocalization Index Manders: Control: 0.5 ± 0.03; Doxorubicin: 0.7 ± 0.03; Etoposide: 0.7 ± 0.01) (**Figure 1E**). Considering the confocal microscopy’s limited resolution to 250 nm, and the fact that the ER-mitochondria distance associated with MERCS falls within a range of 10 to 50 nm, we opted for Transmission Electron Microscopy (TEM) analysis, acknowledged as the gold standard technique, to assess the percentage of MERCS coverage and the distance between ER and mitochondria (**Figure 1F**). Similarly, as shown with confocal microscopy, both Doxo and Eto-induced TIS shows and increase percentage of MERCS coverage, with respect to growing cells (CT: 19.3 ± 1.9%; Doxo: 35.3 ± 2.7%; Eto: 45.6 ± 2.3%). Interestingly, the distance between ER and mitochondria shows a small but consistent increase in both Doxo and Eto-induced TIS with respect to growing cells (CT: 3 0, 13 ± 0.9 nm; Doxo: 33, 25 ± 1.0 nm; Eto: 33, 25 ± 0.8 nm).

**Figure 1.**
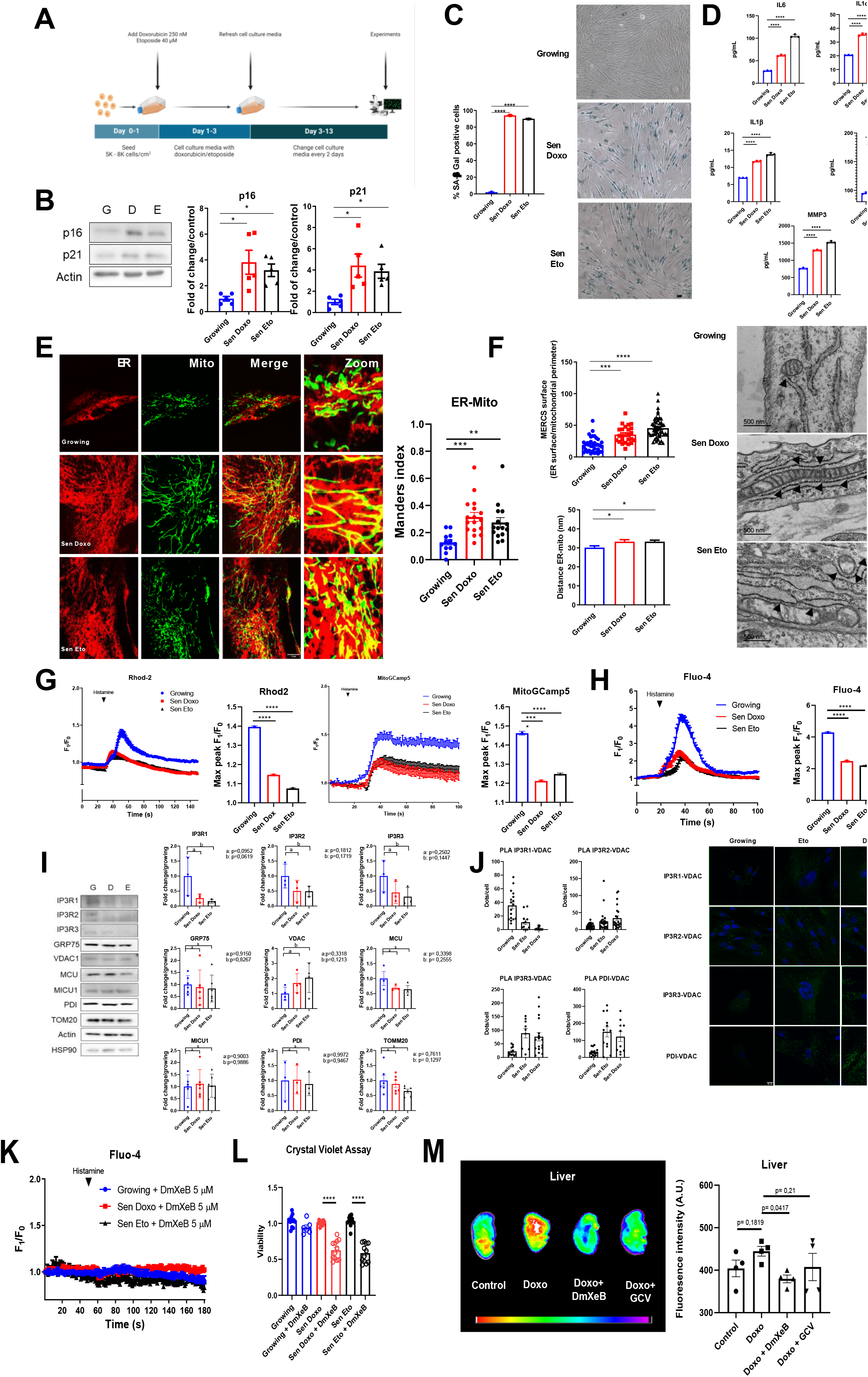
Therapy induces senescence (TIS) increase MERCS, but decreases the ER-mitochondria Ca^2+^ flux, which turns essential for TIS cell survival. **A)** Schematic diagram of induction and establishment of therapy-induced senescence. **B)** Representative Western blot of p16 and p21 in growing and senescence cells induce with doxorubicin (Sen Doxo) and etoposide (Sen Eto). Bar graph: p16/actin and p21/actin expressed as average fold change over basal levels (growing cells). N=5, Mean ± S.E., *p<0.05 (ANOVA Test) **C)** Representative images of SA-β-gal staining in growing, Sen Doxo and Sen Eto and quantitative analysis of stained cells (100 cells per N, N=3; ****p≤0.0001, ANOVA test). Bar: 50 μm. **D)** Concentration of IL6, IL-1α, IL-1β, MMP3, CXCL1, determined by Luminex assay in supernatant of growing, Sen Doxo and Sen Eto cells. (N=3, ****p≤0.0001, ANOVA Test). **E)** Representative images of growing, Sen Doxo and Sen Eto labeled with KDEL-BFP to visualize ER and 100 nM TMRE to visualize mitochondria. Bar: 10 μm. Bar graph: Manders index as an indicator of colocalization (20 – 30 cells per N; N=3; **p≤0.01 *** p≤0.005, ANOVA test). **F)** Representative TEM images and quantification of contact surface area between ER and mitochondria and the distance between ER and mitochondria in growing sen Doxo and Sen Eto. Arrow heads show MERCS (N=3, 50-70 mitochondria per N, *p<0.05, ANOVA test). **G)** Representative traces of mitochondrial Ca^2+^ responds in growing, Sen Doxo and Sen Eto cells challenge with histamine using Rhod-2AM and MitoGCamp5. Bar graphs: Quantification of peak Rhod-2AM (N=3. Mean ± SEM. 50 cells were analyzed in each experiment) and MitoGCamp5 (N=3. mean ± SEM. 20 cells were analyzed in each experiment) fluorescence. ****p≤0.0001, ANOVA test). **H)** Representative traces of cytoplasmic Ca^2+^ signals in growing, Sen Doxo and Sen Eto cells challenge with histamine using Fluo-4AM. Bar graph: Quantification of peak Fluo-4AM fluorescence (N=3. Mean ± SEM. 50 cells were analyzed in each experiment. ****p≤0.0001 ANOVA test). **I)** Representative Western blot of MERCS proteins in growing, Sen Doxo, and Sen Eto. Bar graphs are expressed as average fold change over basal levels (growing cells). N=3-6, mean ± S.E. (ANOVA test). **J)** Representative images of Proximity Ligation Assay (PLA) of the 3 IP3R isoforms with VDAC1, and ER marker PDI with VDAC1 in growing, Sen Doxo and Sen Eto cells. Bar graphs: Quantitative analysis of number of dots per cells. **K)** Representative traces of cytoplasmic Ca^2+^ signals using Fluo-4AM in growing, Sen Doxo and Sen Eto cells treated with 5 μM DmXeB for 1h and challenge with histamine. (N=3. Mean ± SEM. 10-15 cells were analyzed in each experiment). **L)**Crystal Violet assay to evaluate cellular viability. (N=3-4. ****p≤0.0001. ANOVA test). **M)** Representative fluorescence images of p16-3MR mice liver treated with two rounds of Doxo (30mg/k) followed by four rounds of either DmXeB (30mg/k) or ganciclovir. Bar graph: Quantitative analysis of fluorescence intensity (N=4. Means ± SEM. Control: 404.2 ± 19.6, Doxo: 444.6 ± 11.6, Doxo + DmXeB: 379.5 ± 9.09 and Doxo + GCV: 407.5 ± 32.0 A.U. ANOVA test).

To assess the Ca^2+^ flux between the ER and mitochondria, we evaluated mitochondrial Ca^2+^ uptake using two approaches: Rhod-2AM and the protein sensor MitoGCamp5. Surprisingly, both TIS cell models exhibited decreased mitochondrial Ca^2+^ uptake following stimulation with histamine, an inducer of Ca^2+^ release mediated by IP3R (**Figure 1G**). Measurement of cytosolic Ca^2+^ using Fluo-4AM revealed a concurrent decrease in IP3R-mediated Ca^2+^ release from the ER in both TIS cell models (**Figure 1H**).

To unravel the underlying mechanisms responsible of the reduced ER Ca^2+^ release and transfer to the mitochondria, we scrutinized the levels of well-characterized MERCS resident proteins, particularly those implicated with ER-mitochondria Ca^2+^ flux. Intriguingly, we observed decreased levels of the three IP3R isoforms in both TIS cell models compared to growing cells, while other resident proteins such as GRP75, VDAC, MCU, MICU1, PDI, and TOMM20 did not show statistically significant changes (**Figure 1I**). Furthermore, employing Proximity Ligation Assay (PLA), we detected a reduced proximity between IP3R and VDAC (PLA IP3R1-VDAC) in both TIS cell models compared to control growing cells. Intriguingly, the PLA assay revealed an increase in proximity between the ER resident protein PDI and VDAC (PLA PDI-VDAC) (**Figure 1J**). Altogether, our findings indicate that both TIS cell models are characterized by an increase in structural interactions between ER and mitochondria. Notably, these interactions specifically exclude type 1 IP3R isoform, resulting in a reduced Ca^2+^ flux between the ER and mitochondria.

Our group has previously demonstrated the critical role of the Ca^2+^ flux between the ER and mitochondria is crucial in cell survival. Given our observation of a decrease in Ca^2+^ flux between the ER and mitochondria, we aimed to assess the essentiality of this Ca^2+^ flux becomes for TIS cells. To this end, we used Desmethyl XeB (DmXeB), a competitive and selective inhibitor of IP3R, which has been reported to affect the viability of tumor cells[9]. After confirming that DmXeB inhibits Ca^2+^ release from the ER in both growing and TIS cells (**Figure 1K**), we proceeded to evaluate its effect on cell viability. As seen in **Figure 1L**, DmXeB does not affect the viability of growing cells, but it does affect the viability of Doxo and Eto-induced TIS cells (**Figure 1L**). Finally, to investigate whether DmXeB may exert a senolytic effect *in vivo*, we utilized 20-month-old p16-3MR transgenic mice, which allow for the visualization and elimination of senescent cells upon treatment with Ganciclovir[10]. TIS was induced by 2 rounds of Doxorubicin followed by either 4 rounds of DmXeB or Ganciclovir. Fluorescence intensity associated with the senescence marker p16 significantly decreased in the livers of mice treated with DmXeB (**Figure 1M**). Taken together, these results suggest that reducing Ca^2+^ flux between the endoplasmic reticulum and mitochondria is a target for senolysis in the contexts of TIS, and DmXeB emerges as a promising senolytic agent.

MERCS have been recognized as crucial communication platforms orchestrating various cellular processes, including metabolism and apoptosis[4]. In the context of oncogene-induced (OIS) and oxidative stress-induced senescence (OSIS), studies by some groups demonstrated an increase in MERCS interaction surfaces, concomitant with elevated Ca^2+^ flux between the ER and mitochondria, that served as triggering factor for senescence ([5–8]). However, the involvement of MERCS in TIS remains unexplored to date.

Here, we unveil that in TIS, MERCS display an increase as observed in previous studies; nevertheless, the flux of Ca^2+^ between the ER and mitochondria is diminished. This discrepancy may stem from differences in the nature of the senescence inducers, the time frame post-treatment (12 days in our case), and/or the 3% oxygen concentration utilized for cell culture in this study. These oxygen concentrations have been described as normoxic in tissues[11], in addition to show a decrease in the impact of replicative senescence associated with 21% oxygen[12,13]. Nonetheless, the observed reduction in Ca^2+^ flux during TIS is logical, as it may function as an additional antiapoptotic mechanism.

We acknowledge that our study is limitied to IMR-90 cells, and further investigations are warranted in other mesenchymal and epithelial cell types. Additionally, while our findings are pertain to topoisomerase inhibitors, it remains unknown whether similar outcomes can be replicated with different classes of chemotherapeutic agents that induce senescence. Lastly, comprehensive experiments are required to validate the effects of DmXeB in vivo and its potential as a senolytic agent.

In conclusion, our data demonstrate that TIS cells exhibit increased MERCS and reduced Ca^2+^ flux to the mitochondria, which are crucial for maintaining the homeostatic state of these cells.

Data, materials, methods and supporting data is available from the authors upon request.

## CONTRIBUTIONS

U.A-C and C.C., designed and wrote the manuscript. U.A-C performed Ca^2+^ experiments. A.P-H, C.Q and G.B., performed confocal microscopy experiments and Western blotting. E.S-P, P.M-C, and C.C-C, performed cytometry assay. O.D-R performed plasmid extraction. J.A.L-D, S.R-L, and J.M.V. performed TEM experiments. H.H, G.Q, C.G-S, S.L., E.S-H, and M.J.K. conducted mice experiments. H.H. performed cytokine measurements. A.L., C.G-B, J.M., and A.Z. contributed to interpretation of the results and provided feedback. All authors read and approved the final manuscript.

## ACKNOWLEDGEMENTS

We thank Judith Campisi and Pierre-Yves Desprez from the Buck Institute for Research on Aging for their mentorship and for guiding us in establishing TIS models.

